# Malignant Schwann cell precursors mediate intratumoral plasticity in human neuroblastoma

**DOI:** 10.1101/2020.05.04.077057

**Authors:** Thale K. Olsen, Jörg Otte, Shenglin Mei, Polina Kameneva, Åsa Björklund, Emil Kryukov, Ziyi Hou, Anna Johansson, Erik Sundström, Tommy Martinsson, Susanne Fransson, John Inge Johnsen, Per Kogner, Igor Adameyko, Peter V. Kharchenko, Ninib Baryawno

## Abstract

Neuroblastoma is a heterogeneous embryonal malignancy and the most deadly tumor of childhood, although a minor subset may show spontaneous differentiation. It arises from the multipotent neural crest lineage during development. Some of this multipotency is retained in neuroblastoma, which can give rise to both adrenergic and mesenchymal tumor cells. The mechanisms enabling such dual fates are unknown, but likely help neuroblastoma to evade existing therapies. To understand neuroblastoma plasticity, we analyzed patient tumors using single-cell transcriptomics. In addition to the heterogeneous adrenergic and mesenchymal populations, we identify a subpopulation of malignant cells resembling Schwann cell precursors (SCPs). This SCP-like population connects the adrenergic and mesenchymal compartments through transitions structurally reminiscent of the SCP cell-fate decision fork that occurs during normal development. While the directionality of such transitions in neuroblastoma remains to be established, this finding expands the potential reservoirs of malignant cells, and suggests intratumoral plasticity mechanisms relevant for therapeutic resistance and relapse.

---

Neuroblastoma is a pediatric extra-cranial neuroendocrine cancer, believed to arise early at prenatal stages from the sympatho-adrenal lineage of the neural crest ^1,2^. It is a complex disease that can result in diverse clinical outcomes. In some patients, the tumors regress spontaneously. Others respond well to existing treatments. But for the high-risk group, which constitutes approximately 40% of all cases, the prognosis remains dire despite collaborative efforts in basic and clinical research. ^1,3,4^ The disease is also heterogeneous on the cellular level. Examination of neuroblastoma cell lines and patient samples has revealed that neuroblastoma can give rise to two distinct tumor cell subtypes: adrenergic and mesenchymal ^5,6^. The two subtypes can occur within the same tumor, with the mesenchymal cells appearing to be more resistant to chemotherapy ^5^. The ability of the two lineages to interconvert may contribute further to therapeutic resistance. *In vitro* studies of cell lines suggest that adrenergic-to-mesenchymal transitions may be enabled by activation of the NOTCH signaling pathway which is also involved in cell identity choices in development ^7^. Despite the clinical relevance, however, the mechanisms underlying plasticity of neuroblastoma tumors remain poorly understood.

Many tumors are known to activate and replay developmental mechanisms to attain plasticity that ultimately promotes resistance to therapy ^8^. The neural crest lineage, from which neuroblastoma originates, is remarkably flexible, and can give rise to both mesenchymal and adrenergic lineages during normal development ^9–13^. In particular, using lineage tracing experiments in mice, we have recently observed that these adrenergic and mesenchymal cell types share an immediate developmental progenitor, represented by neural crest cells diverging at last towards sympatho-adrenal and mesenchymal sub-lineages ^14^. Furthermore, this bifurcation involved transient, bipotent progenitors which initially activate both adrenergic and mesenchymal transcriptional programs. The precise developmental origin of neuroblastoma, however, remains unclear, and likely involves downstream neural crest lineages. In an embryo, neural crest cells settle on the growing nerves and transform into Schwann cell precursors (SCPs), which express the most archetypical part of the neural crest signature and retain neural crest-like multipotency ^9,15^. SCPs are retained for a considerable time during development and continuously give rise to chromaffin cells and some sympathetic neurons of the adrenal gland region which, in turn, is the most common anatomical location of neuroblastoma ^9,10,16^. Here, by performing in-depth profiling of human neuroblastoma tumor samples at a single-cell level, we show continuous transitions between the two previously identified mesenchymal and adrenergic neuroblastoma subpopulations. Surprisingly, this transition appears to be mediated by a third cell state, which mimics both molecular and phenotypic features of normal SCPs.

To understand what molecular mechanisms may underlie neuroblastoma plasticity, we used single-cell transcriptomics and genotyping arrays to characterize 17 neuroblastoma samples across different clinical risk groups, ages, and genetic subsets (**Fig. 1A** and table S1). Fresh samples, obtained from surgical resections and core needle biopsies, were used to generate single-cell suspensions and assayed using single-cell RNA-seq (scRNA-seq, 10x Genomics Chromium), yielding a total of 72,606 high quality cells (see Methods). To overcome inter-individual variation, the samples were integrated using a graph-based *Conos* approach ^17^ (**Fig. 1B**), which revealed at least 18 major cell subpopulations common to most samples, spanning tumor, immune and stromal compartments (**Fig. 1, B and C**, and fig. S1, A to D). Among the mesenchymal stromal populations we observed well-defined endothelial cells, pericytes and myofibroblasts. The remaining, connected populations expressed mesenchymal (*PRRX1, LEPR, PDGFRA, DCN*) and adrenergic (*TH, DBH*) genes (**Fig. 1D**), indicative of mesenchymal and adrenergic tumor origin, respectively ^5,6^. Unexpectedly, these two putative tumor populations were connected by a bridge that could not be attributed to capture of cell doublets (fig. S1E) and showed expression of key neural crest and Schwann lineage markers (*SOX10, S100B*, **Fig. 1, C and D**, and fig. S1, C and D).

**Figure 1.**
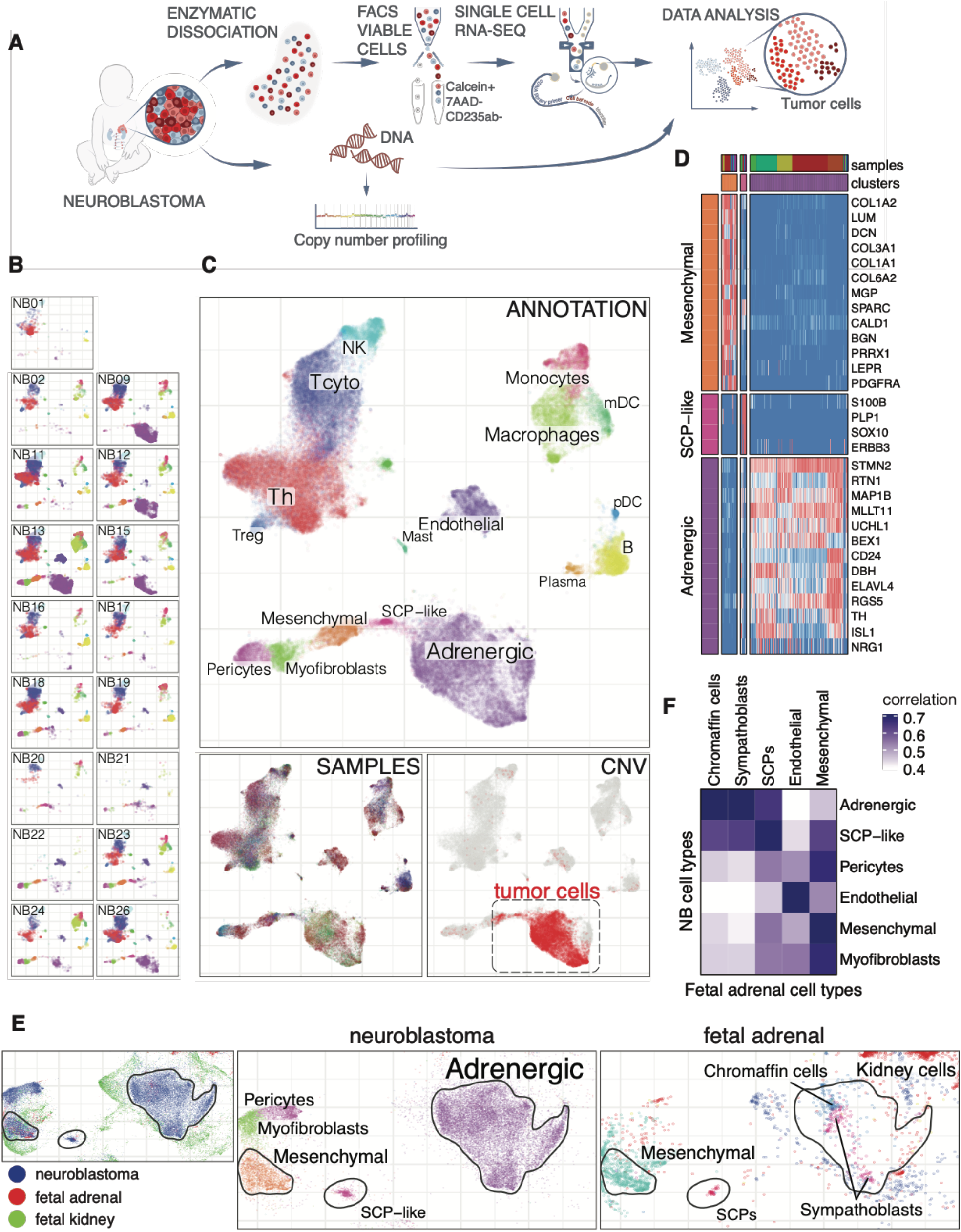
Single-cell RNA sequencing of human neuroblastoma reveals a third, distinct population of SCP-like bridge cells resembling normal fetal SCPs. (**A**) Illustration of study design and experimental procedures. (**B** and **C**) Integrative analysis of scRNA-seq samples from 17 neuroblastoma samples, visualized using a common UMAP embedding. B. Individual samples. C (top) - major cell populations. Treg: regulatory T cells, Th: T helper cells. Tcyto: cytotoxic T cells, mDC: myeloid dendritic cells, pDC: plasmacytoid dendritic cells. C (bottom left) - distribution of cells from individual samples. C (bottom right) - red color shows malignant cells, as determined based on the elevated (5% FDR) average expression levels within the regions of patient-specific genomic gain. (**D**) Expression of select marker genes in Adrenergic, Mesenchymal and SCP-like populations. (**E**) Joint alignment of neuroblastoma, fetal adrenal and fetal kidney scRNA-seq datasets is shown, colored by data resource (left), neuroblastoma cell annotation (middle), and human adrenal data annotation (right). Solid contours show positions of the neuroblastoma Adrenergic, Mesenchymal and SCP-like populations. (**F**) Correlation of the overall expression profiles between different neuroblastoma and fetal adrenal cell types illustrates best matching cell populations.

To distinguish malignant cells from normal, we evaluated systematic up- or down-regulation of genes within the cancer-specific copy number variants (CNVs) ^18^. Large-scale genomic amplifications, such as 2p gain, should on average lead to increased expression of the genes within them, and deletions to average expression reduction ^19^. Using CNV profiles from array-CGH data from the same tumors (fig. S1F), we derived a combined score that quantifies the extent to which the expression pattern of a given cell reflects the CNV profile of a given patient. Indeed, we found that the high-scoring cells (above 5% FDR threshold, see Methods) comprise the clusters carrying adrenergic, mesenchymal and neural crest signatures (**Fig. 1C**, and fig. S1, G and H).

Neuroblastomas are thought to arise during differentiation of neural crest-derived cells into the adrenal lineages ^2^. To understand how the observed malignant populations are related to normal adrenal tissue, we performed single-cell transcriptional profiling of adrenal tissue from a week 8 human fetus (fig. S1J). Integration of this fetal adrenal data with neuroblastoma samples and recently published fetal kidney measurements ^20^ confirmed close resemblance of normal developmental mesenchymal and adrenergic (chromaffin, sympathoblasts) cells to the distinct subpopulations in neuroblastoma as demonstrated by earlier studies ^5,6^ **(Fig. 1, E and F**, and fig. S1K). Unexpectedly, the malignant bridge population aligned with SCPs normally observed in fetal development **(Fig. 1, E and F**, and fig. S1K), which suggested the existence of a third distinct neuroblastoma state mimicking human fetal SCPs.

The SCP-like bridge population expressed a broad range of markers characteristic of Schwann cell lineage ^9^, including *S100B, SOX10, ERBB3, FOXD3* and *PLP1*. Numerous cells within this cluster also expressed genes related to myelination (e.g. *LGI4, PMP2, MPZ*) and axon guidance (*SEMA3B*) typical of more mature myelinating Schwann cells (**Fig. 2, A and B**, and fig. S1C). This suggests the existence of a diversity reflecting the spectrum of Schwann cell lineage phenotypes ranging from less mature SCPs to mature myelinating cells in the tumors. The similarities between normal SCPs of the developing fetus and SCP-like neuroblastoma cells extended to important functional traits. For instance, using SOX10 as a marker of SCPs, we performed immunostainings to validate the expression and spatial context of SOX10+ cells in neuroblastoma samples. SOX10+ cells were detected at varying frequencies in all stained samples (**Fig. 2, C and D**, and fig. S2, A and B). The migration and differentiation of normal SCPs during development is guided by nerve fibers ^21,22^. The association with the nerves is maintained through the ERBB3 receptor, which in turn binds NRG1 – a critical signal secreted by the nerves ^23,24^. Similar to the SCPs of the developing embryo, the vast majority of SOX10+ cells in neuroblastoma were positioned close to NF200+ neurofilaments (**Fig. 2, C and D**). SOX10+ cells expressed ERBB3 and the ligand NRG1 was expressed adjacent to SOX10+ cells, indicating that the crucial SCP niche is preserved in neuroblastoma (**Fig. 2E** and fig. S2C). Some of the neurofilaments appear to be recruited nerve bundles, while others appear diffuse and are likely produced by the adrenergic subpopulation of the tumor cells which also expresses NRG1 **(Fig. 1D).** Although the SCP-like neuroblastoma cells express many glial genes, with some cells being myelin protein zero (MPZ) positive, most SOX10^+^ cells do not express the MPZ protein (fig. S2D), suggesting that they do not represent a mature Schwann cell phenotype.

**Figure 2.**
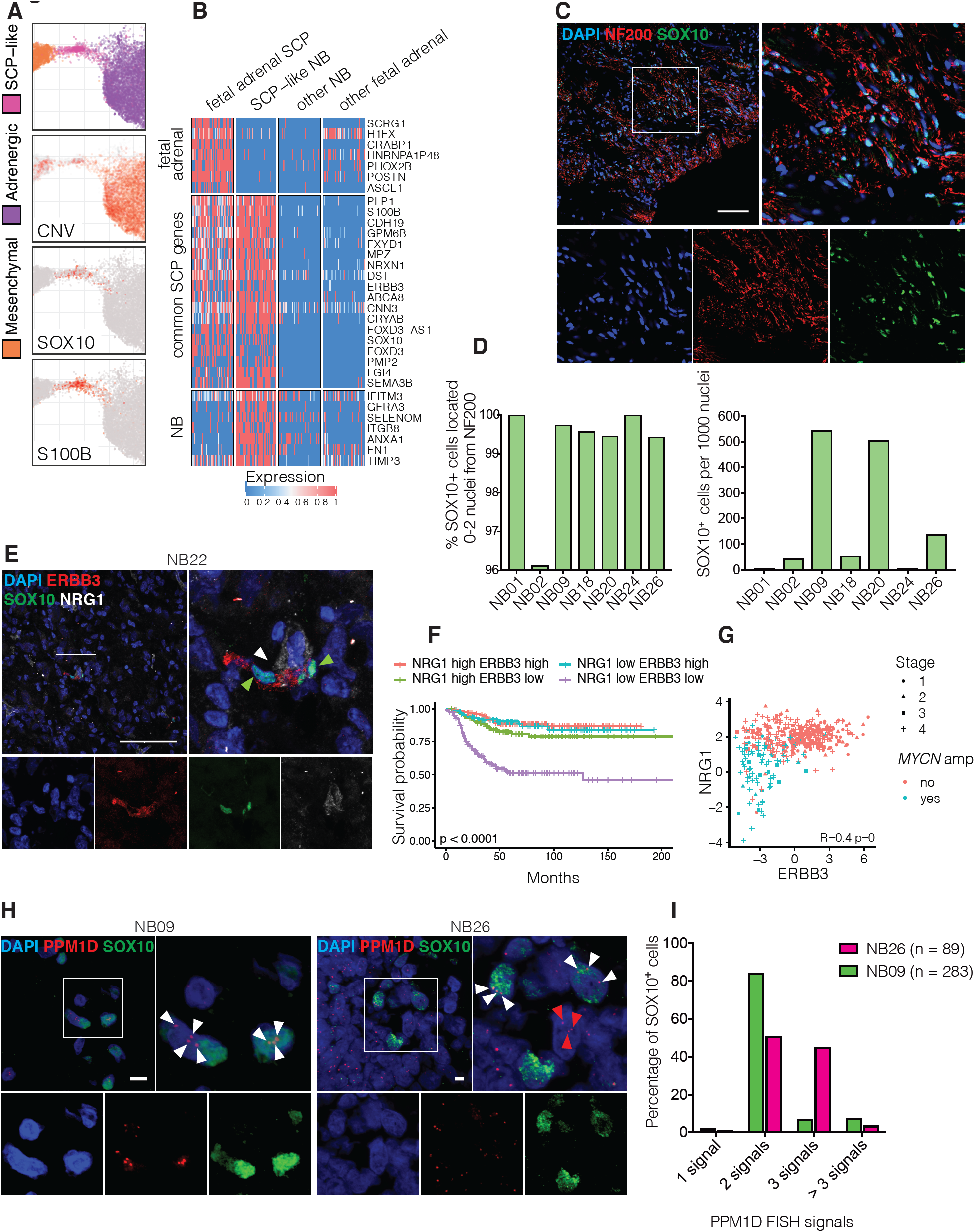
Schwann cell precursor-like cells in neuroblastoma are malignant and retain key developmental niche traits. (**A**) Enlarged view of the region connecting adrenergic and mesenchymal cell populations in the joint embedding of the neuroblastoma samples, colored by cell annotation (top), malignant cells based on the copy number variation expression scores (CNV), and key SCP marker genes. (**B**) Common and discordant markers of the SCP population found in fetal adrenal data (first column) and the SCP-like neuroblastoma population (second column) are shown as a heatmap, separating feta adrenal-specific (top), common (middle), and neuroblastoma-specific (bottom) markers. (**C**) Immunohistochemistry for neurofilament 200 (NF200; shown in red) and SOX10 (shown in green) on a section from sample NB20. SOX10+ nuclei are situated in close proximity to NF200 protein. Scale bar indicates 100 μm. (**D**) Quantification of SOX10+ cells and their relation to NF200 in seven neuroblastoma samples. Left panel: Percentage of SOX10+ cells located at a distance of 0-2 nuclei from NF200 protein. Right panel: Number of SOX10+ cells per 1000 DAPI+ nuclei. Total number of DAPI+ nuclei ranged from 2518 (NB09) to 11002 (NB26). Cells on 3-7 scans were included in the calculation. (**E**) Immunohistochemistry for SOX10 (green), the receptor tyrosine kinase erbB-3 (ERBB3; shown in red), and its ligand neuregulin-1 (NRG1, shown in white). Green arrows indicate SOX10+ cells co-expressing ERBB3. White arrows indicate presence of NRG1. Scale bar indicates 100 μm. (**F**) Kaplan–Meier survival curves for neuroblastoma patients (Su et al. dataset, reference no. 26; INSS stage 4S patients excluded, n = 445) stratified into four groups based on *ERBB3* and *NRG1* expression. (**G**) The relationship between *ERBB3* and *NRG1* expression in bulk RNA-seq data is shown as a scatter plot. Each point corresponds to a patient (Su et al. dataset, INSS stage 4S patients excluded, n = 445), with color designating *MYCN* amplification status, and shape designating INSS stage. Expression values are shown as log FPKM. (**H)** Validation of SOX10+ cell malignancy using combined FISH and immunohistochemistry in two samples. SOX10 shown in green, *PPM1D* FISH signal shown in red. White arrows indicate aberrant signal in SOX10+ cells, red arrows indicate aberrant signal in SOX10-cells. Scale bar indicates 10 μm. (**I**) Quantification of *PPM1D* FISH signal in SOX10+ nuclei in samples NB09 and NB26. Total number of DAPI+ nuclei evaluated: n = 628 (NB09); n = 606 (NB26). Legend indicates total number of SOX10+ cells evaluated in each sample.

The neuroblastoma SCP-like state and the normal fetal adrenal SCPs also displayed notable differences. Whereas the fetal SCPs obtained from the adrenal gland expressed *PHOX2B* and *POSTN*, inclining them towards differentiation into the mature sympathoadrenal lineage, very few tumor SCP-like cells expressed these genes (**Fig. 2B**). Instead, the SCP-like tumor cells expressed genes such as the artemin receptor *GFRA3*, recently shown to promote hepatocellular carcinoma initiation and growth ^25^, and the integrin subunit *ITGB8*, which is implicated in differentiation and radiosensitization of glioblastoma ^26^ (**Fig. 2B**). As expression of oncogenes suggested that SCP-like cells may enhance tumor growth, we examined the association of key SCP genes with disease progression. The analysis of gene expression data from larger patient cohorts ^27,28^, however, demonstrated that the presence of key SCP-like genes, exemplified by *SOX10*, was instead associated with prolonged patient survival (fig. S2, E and F). This association was robust with respect to the choice of the SCP-like signature genes, and even extended to the NRG1 ligand which is not expressed by the SCP-like cells themselves but is required by normal SCPs (**Fig. 1D and Fig. 2F**). Much of the difference in survival, however, could be attributed to the lack of SCP-like signature in the patients with amplifications of *MYCN* (**Fig. 2G** and fig. S2, G and H), suggesting distinct etiology in these tumors.

In contrast to malignant SCP-like cells suggested by the results above, the occurrence of stromal Schwann populations in neuroblastic tumors is well established ^29^. They are thought to be recruited to the tumor from surrounding non-malignant tissue ^30^, and their abundance is used to classify tumor subtypes ^31^. Cases with a high proportion of Schwann cell stroma, such as differentiated ganglioneuromas and stroma-rich ganglioneuroblastomas, are less aggressive and have better clinical outcomes ^3,31^. This observation is consistent with the survival differences noted above (fig. S2, E and F). Our observations are, instead, aligned with a handful of reports arguing that the tumors can produce malignant Schwann lineage directly ^32–34^. While the expression-magnitude and allele-based inference of genetic aberrations in the scRNA-seq data strongly indicates that SCP-like cells carry cancer-specific CNVs (**Fig. 2A** and fig. S1, G and H), we sought to validate these findings using an orthogonal technique. Specifically, we combined fluorescence in situ DNA hybridization (FISH) targeting 17q gain (*PPM1D* locus) with immunofluorescence stainings for SOX10, the adrenergic marker ISL1 and the mesenchymal marker PDGFRA (**Fig. 1D**). The samples selected for staining all displayed 17q gain as evaluated by array CGH (fig. S1F). SOX10 immunofluorescence and FISH stainings were of sufficient quality for evaluation in two samples: NB09 and NB26. Similar to adrenergic ISL1+ cells harboring an aberrant number of *PPM1D* signals (fig. S2I), we found that a fraction of the SOX10+ SCP-like cells exhibited an increased and abnormal number of intra-nuclear signals hybridizing to the *PPM1D* locus (**Fig. 2, H and I)**. Such shared genomic abnormalities between the malignant adrenergic and SCP-like cells confirm that SCP-like cells can be produced by the tumor, and that the SCP-like population likely represents a mixture of malignant and recruited cells.

In our joint analysis of neuroblastoma samples, the SCP-like population was connected to the adrenergic and mesenchymal tumor cell lineages by continuous bridges (**Fig. 1C**), reflecting a gradual change of expression characteristic of a continuous differentiation process (**Fig. 3A**). Such lineage arrangement is analogous to a neural crest cell fate fork that we have recently described in normal murine development ^14^. In that context, the migrating neural crest cells, which are immediate progenitors of the SCPs, similarly give rise to both autonomic neurons (expressing *Phox2b, Ascl1, etc.*) and mesenchymal populations (*Prrx1, Fli1*) with a mixed phenotype identified at the fate split point (*Prrx1, Phox2b, Sox10*) (**Fig. 3, B and C**). The SCPs themselves largely resemble neural crest and are known to produce both adrenergic ^9,10^ and mesenchymal lineages ^11–13^ during development, though the detailed transcriptional trajectories of such transitions have not been characterized in either human or mouse. Despite the detected tumor-specific differences, the observed homology of lineage arrangements suggests that intratumoral plasticity of neuroblastoma may involve normal developmental mechanisms.

**Figure 3.**
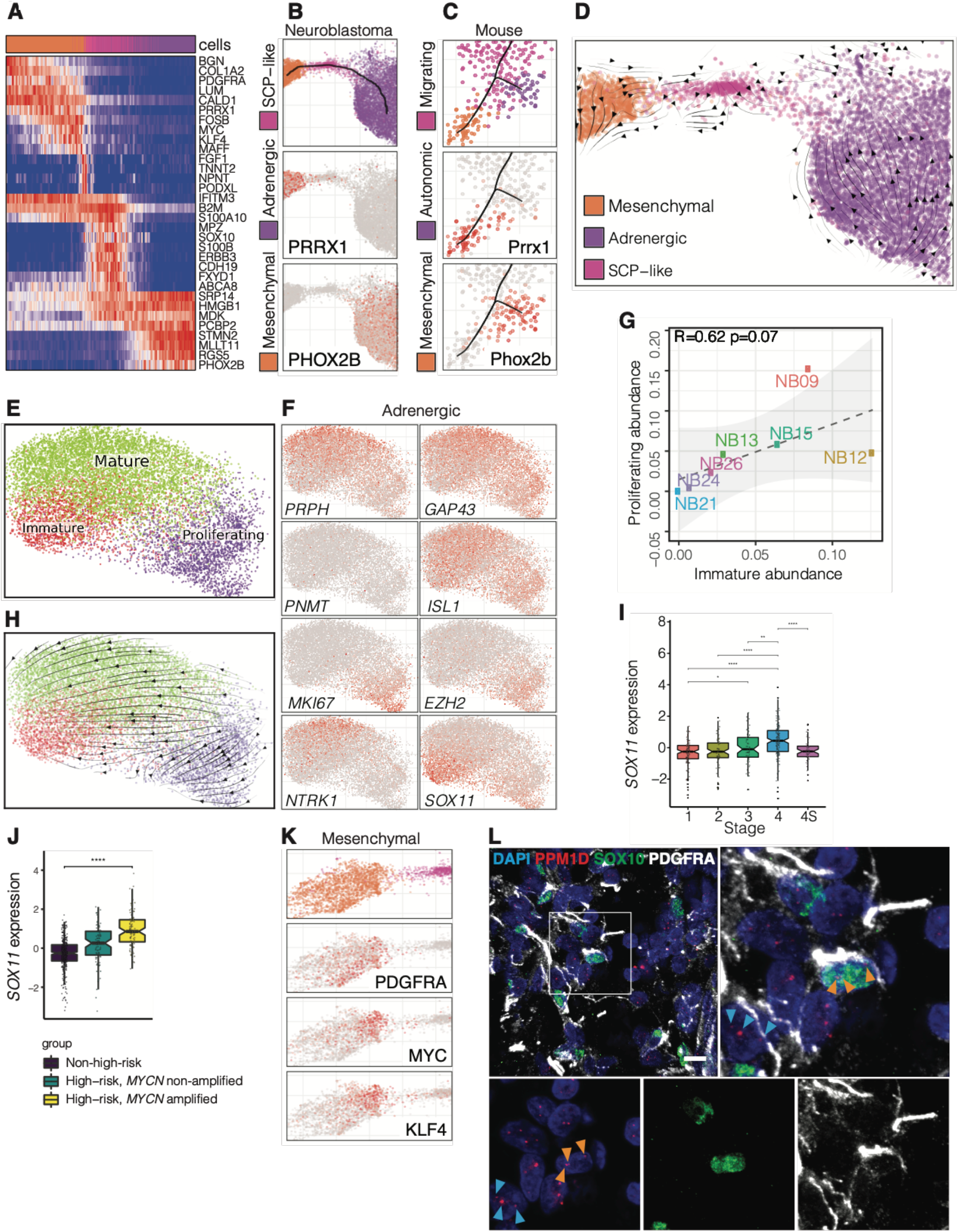
Intratumoral plasticity in human neuroblastoma suggest transitions within and between neuroblastoma cell states. (**A**) The heatmap shows gene expression transitions within the bridge region (Fig. 2A) connecting mesenchymal, SCP-like, and adrenergic populations. The cells were ordered by pseudotime shown in panel B. (**B**) Pseudotime within the bridge region (top) is shown along with expression of *PRRX1* and *PHOX2B*; markers of the mesenchymal and adrenergic populations, respectively. (**C**) Analogous developmental cell fate decision fork in mouse neural crest, separating mesenchymal (*Prrx1*) and autonomic (*Phox2b*) branches. (**D**) RNA velocity analysis of the transitions surrounding SCP-like neuroblastoma state, estimated on combined data from all samples. (**E**) Separate re-analysis of the neuroblastoma adrenergic population shows separation of three major subpopulations. (**F**) Expression of markers in the re-analyzed adrenergic embedding (E). (**G**) Scatter plot illustrates significant correlation between the abundance (assessed as a proportion of all cells in the sample) of the *Immature* (x axis) and *Proliferating* (y axis) adrenergic subpopulations. (**H**) RNA velocity analysis showing predicted dynamics within the adrenergic compartment. (**I-J**) Expression of *SOX11* in bulk RNA-seq data (Su et al. dataset, all patients, n = 498) of different patients is shown, stratifying patients by INSS stage (3I) or INRG risk stratification (3J). *: p < 0.05. **: p < 0.01. ****: p < 0.0001. Expression values are shown as log FPKM. (**K**) Enlarged view of the mesenchymal population is shown along with expression of select marker genes. (**L**) Validation of PDGFRA+ cell malignancy using combined FISH and immunohistochemistry in sample NB26. *PPM1D* FISH signal shown in red, SOX10 in green, PDGFRA in white. Scale bar indicates 10 μm. Light blue arrows indicate SOX10-, PDGFRA+ cell with aberrant *PPM1D* signal (three dots). Orange arrows indicate SOX10+, PDGFRA+ cell with aberrant *PPM1D* signal (three dots).

Developmental differentiation of SCPs into the adrenal compartment can produce both neuroendocrine chromaffin and sympathetic neurons ^9,10^. The adrenergic cells of neuroblastoma generally resembled sympathoblasts, expressing *PRPH* and *GAP43*, and lacked chromaffin markers such as *PNMT* (**Fig 3F**). To analyze heterogeneity within the adrenergic compartment, we excluded a subcluster of cells that lacked CNV genomic gain signal (**Fig. 1C** and fig. S3A), and re-ran integration analysis (**Fig. 3E**). We found three major clusters of malignant adrenergic cells. The first, *Proliferating* cluster, was marked by a strong mitosis-related signature, exemplified by *MKI67* (**Fig. 3, E and F**, and fig. S3B). Interestingly, the cluster was also distinguished by high expression of the epigenetic modifier *EZH2*. This gene has been previously shown to promote neuroblastoma growth through inhibition of tumor suppressor genes ^35,36^ and inhibition of drivers of sympathoblast differentiation such as *NTRK1* ^37^. The second, *Mature*, cluster showed expression of *NTRK1, PRPH, GAP43* and other markers characteristic of differentiation into sympathetic neurons ^38^, and was also enriched with genes related to the synaptic machinery (**Fig. 3F** and fig. S3, B and C). Notably, *NTRK1* expression in the *Mature* cluster parallels with a good outcome in NTRK1^+^ neuroblastoma. NTRK1 (also known as TrkA) is a landmark of favorable neuroblastoma, in particular the subset of infant stage 4S/MS tumors which often regress spontaneously ^3,39^. Finally, the third *Immature* cluster lacked both maturation and cell cycle markers, but expressed high levels of *SOX11*, a transcription factor important for early phases of pro-adrenergic differentiation ^40^, and *CTNNB1*, a key downstream component of the canonical Wnt signaling pathway (**Fig. 3F** and fig. S3B). Interestingly, *in vitro* CRISPR screens have demonstrated that *SOX11* dependency is a highly distinguishing feature of neuroblastoma cell lines ^41^ (fig. S3D). Consistently, examining larger bulk RNA-seq cohorts, we observed that *SOX11* and other key markers of the *Immature* cluster were associated with poor patient prognosis (**Fig. 3, I and J**, and fig. S3, E and F). As expected, poor outcomes were also linked to *Proliferating* signatures (fig. S3F), whereas markers of the *Mature* population were linked to improved survival. The three populations likely transition into each other. Analysis of RNA velocity ^42^ suggested that the *Proliferating* cluster potentially contributes to the *Immature* and *Mature* subpopulations (**Fig. 3H** and fig. S3G). We noted that despite being expressed by the *Immature* cluster, *SOX11* expression correlated with mitotic markers in bulk RNA-seq data (fig. S3H). Such pattern could arise if the abundance of the *Immature* and *Proliferating* populations was correlated. Indeed, we found that the proportion of the two populations correlates across the examined tumors (**Fig. 3G** and fig. S3I), suggesting that the *Proliferating* cluster actively replenishes the *Immature* population. This resembles the concept of tumor stem cells and differentiation-like dynamics ^43^.

Heterogeneity was also apparent within the mesenchymal compartment. The expression analysis of CNV signatures indicated that the overall mesenchymal cluster represents a mixture of malignant and non-malignant cell populations (**Fig. 1C**). We validated the presence of malignant mesenchymal cells ^5,6^ through combined immunofluorescence staining (PDGFRA) and FISH (*PPM1D* at 17q) and identified SOX10+/PDGFRA+ as well as SOX10-/PDGFRA+ cells with *PPM1D* gain (**Fig. 3L)**, indicating a possible transition between malignant SCP-like and mesenchymal neuroblastoma cell states. Towards the end of the bridge connecting SCP-like cells with the mesenchymal population, we observe expression of a coordinated group of genes (**Fig. 3A** and fig. S3J), including the *FGF1* growth factor, enriched for anatomical and tissue development functions (GO:0048646 Q-value 4.4 × 10^−6^). The mesenchymal population adjacent to the bridge, which is most likely to be malignant (**Fig. 1C**), also exhibited a distinct expression signature with high levels of *MYC, KLF2* and other pluripotency-related factors (**Fig. 3, A and K**). Despite the lack of a mitotic signature, the presence of distinct mesenchymal populations expressing pluripotency and growth signals as described above suggests active contribution of the mesenchymal compartment to malignant tumor growth.

Overall, our results demonstrate that in addition to heterogeneous mesenchymal and adrenergic compartments, neuroblastomas contain a malignant SCP-like population with stem-cell like features, and that continuous transitions likely take place between the SCP-like cells and other neuroblastoma compartments (**Fig. 4**). The directionality of these *in vivo* transitions is challenging to establish. RNA velocity analysis indicates likely transitions from the SCP population towards both mesenchymal and adrenergic compartments, depending on the patient (**Fig. 3D** and fig. S3K). Although neuroblastoma is believed to arise during neural crest and sympathoadrenal development, a neuroblastoma stem cell has not yet been identified. If, indeed, the transitions abide by the developmental direction – proceeding from SCPs – it would imply that the SCP-like state constitutes a malignant stem/progenitor population; possibly the neuroblastoma cell of origin. Transitions in the opposite direction would entail de-differentiation of adrenergic or mesenchymal populations into the SCP-like state and cannot be ruled out. Additional studies with larger sample numbers and orthogonal functional experimental techniques will be needed to resolve these alternatives. Knowledge of the machinery enabling neuroblastoma transitions or the identity of a malignant progenitor population would provide an important basis for advancing biological understanding and development of targeted therapeutic interventions.

**Figure 4.**
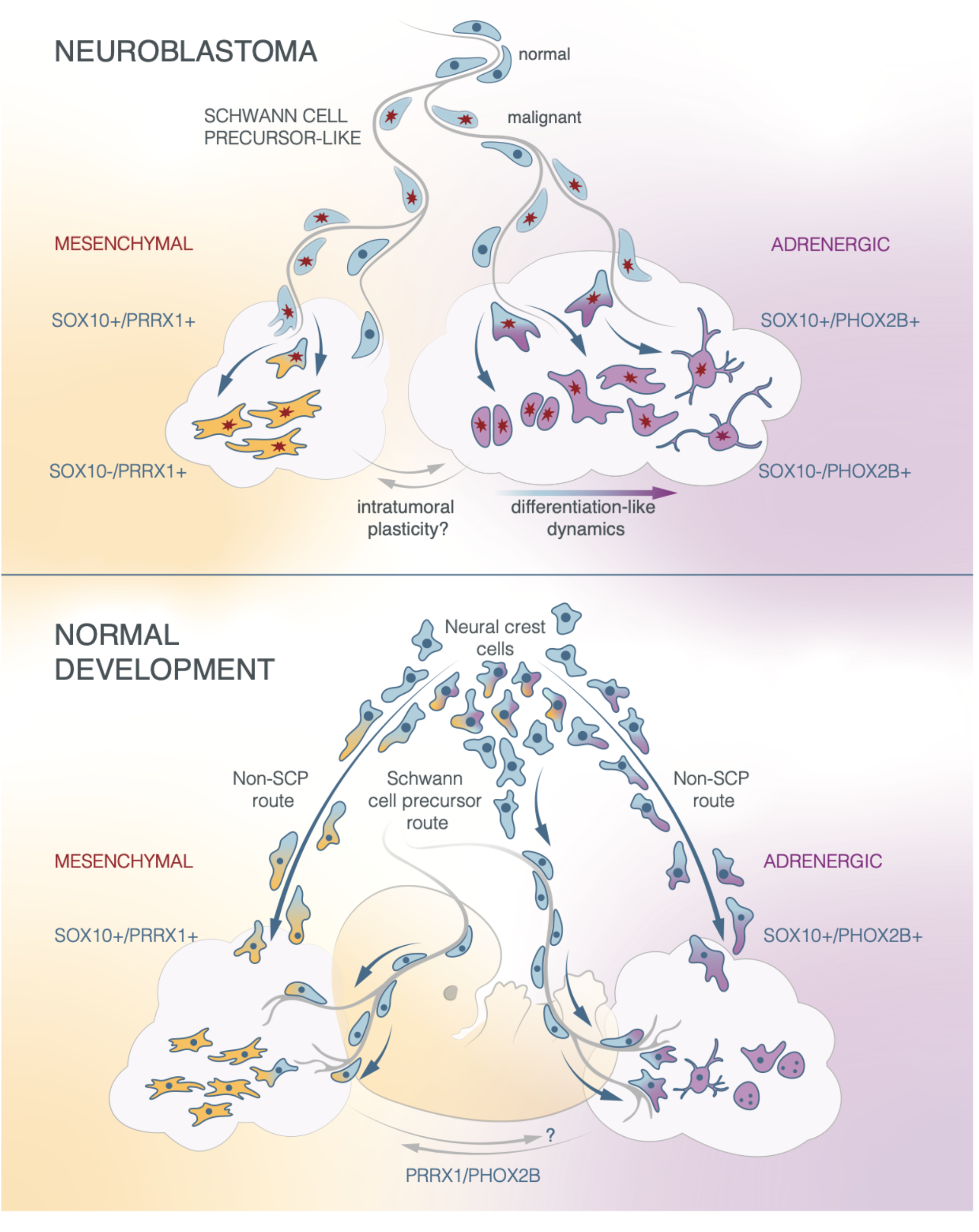
Schematic model summarizing the role of Schwann cell precursors-like cells in neuroblastoma and Schwann cell precursors in normal development. *Upper*: Bipotent, nerve-associated malignant SOX10+ SCP-like cells differentiate into mesenchymal (PRRX1+) or adrenergic (PHOX2B+) tumor cells via gradual transition. Within the adrenergic compartment, subpopulations of various differentiation states exist. This connection of the adrenergic and mesenchymal compartment by SCP-like progenitors resembles the decision fork occurring during normal development. *Lower*: Early neural crest cells which are also progenitors of SCPs initially co-activate adrenergic and mesenchymal transcriptional programs. Cranial neural crest cells also develop into mesenchymal structures of the head, while trunk neural crest cells build sensory and autonomic cell types. Adrenergic chromaffin cells of the adrenal medulla originate to a large extent from SCPs which depend on nerves as their stem cell niche.

## Acknowledgments

We thank Jakob Stenman for help in providing neuroblastoma samples. We thank Bethel Embaie and Ioanna Tsea for reviewing and commenting on the manuscript. We thank Bronte Manouk Verhoeven for aiding immune cell annotations. We thank Olga Kharchenko for help with graphical illustrations.

## Author contributions

Conceptualization, N.B.; Investigation, T.K.O., J.O., S.M., I.A., J.I.J., E.S., P.Ka., P.Ko., P.V.K. and N.B.; Validation, T.K.O., J.O., P.Ka., E.K., Z.H., T.M., S.F.; Computational investigation and analysis, T.K.O., S.M., Å.B., A.J., and P.V.K.; Writing -– Original draft, T.K.O., J.O., I.A., P.V.K. and N.B.; Writing – Review & Editing, T.K.O., E.S., I.A., J.I.J., P.Ko., P.V.K. and N.B.; Funding Acquisition, Resources & Supervision, P. Ko., P.V.K., and N.B.

## Supplementary Materials

Materials and Methods

Figures S1-S3

Data S1-S4

References (43 – 45)

## References

1. Matthay, K. K. et al. Neuroblastoma. Nat. Rev. Dis. Prim. 2, 16078 (2016).

2. Hoehner, J. C. et al. A developmental model of neuroblastoma: Differentiating stroma-poor tumors’ progress along an extra-adrenal chromaffin lineage. Lab. Investig. 75, 659–675 (1996).

3. Cohn, S. L. et al. The International Neuroblastoma Risk Group (INRG) classification system: an INRG Task Force report. J. Clin. Oncol. 27, 289–97 (2009).

4. Pinto, N. R. et al. Advances in risk classification and treatment strategies for neuroblastoma. Journal of Clinical Oncology vol. 33 3008–3017 (2015).

5. van Groningen, T. et al. Neuroblastoma is composed of two super-enhancer-associated differentiation states. Nat. Genet. 49, 1261–1266 (2017).

6. Boeva, V. et al. Heterogeneity of neuroblastoma cell identity defined by transcriptional circuitries. Nat. Genet. 49, 1408–1413 (2017).

7. van Groningen, T. et al. A NOTCH feed-forward loop drives reprogramming from adrenergic to mesenchymal state in neuroblastoma. Nat. Commun. 10, 1530 (2019).

8. Dravis, C. et al. Epigenetic and transcriptomic profiling of mammary gland development and tumor models disclose regulators of cell state plasticity. Cancer Cell 34, 466-482.e6 (2018).

9. Furlan, A. et al. Multipotent peripheral glial cells generate neuroendocrine cells of the adrenal medulla. Science 357, eaal3753 (2017).

10. Kastriti, M. E. et al. Schwann cell precursors generate the majority of chromaffin cells in zuckerkandl organ and some sympathetic neurons in paraganglia. Front. Mol. Neurosci. 12, 6 (2019).

11. Xie, M. et al. Schwann cell precursors contribute to skeletal formation during embryonic development in mice and zebrafish. Proc. Natl. Acad. Sci. U. S. A. 116, 15068–15073 (2019).

12. Joseph, N. M. et al. Neural crest stem cells undergo multilineage differentiation in developing peripheral nerves to generate endoneurial fibroblasts in addition to Schwann cells. Development 131, 5599–612 (2004).

13. Carr, M. J. et al. Mesenchymal precursor cells in adult nerves contribute to mammalian tissue repair and regeneration. Cell Stem Cell 24, 240-256.e9 (2019).

14. Soldatov, R. et al. Spatiotemporal structure of cell fate decisions in murine neural crest. Science 364, eaas9536 (2019).

15. Adameyko, I. et al. Schwann cell precursors from nerve innervation are a cellular origin of melanocytes in skin. Cell 139, 366–379 (2009).

16. Vo, K. T. et al. Clinical, biologic, and prognostic differences on the basis of primary tumor site in neuroblastoma: A report from the International Neuroblastoma Risk Group project. J. Clin. Oncol. 32, 3169–3176 (2014).

17. Barkas, N. et al. Joint analysis of heterogeneous single-cell RNA-seq dataset collections. Nat. Methods 16, 695–698 (2019).

18. Caren, H. et al. High-risk neuroblastoma tumors with 11q-deletion display a poor prognostic, chromosome instability phenotype with later onset. Proc. Natl. Acad. Sci. 107, 4323–4328 (2010).

19. Patel, A. P. et al. Single-cell RNA-seq highlights intratumoral heterogeneity in primary glioblastoma. Science 344, 1396–401 (2014).

20. Hochane, M. et al. Single-cell transcriptomics reveals gene expression dynamics of human fetal kidney development. PLoS Biol. 17, e3000152 (2019).

21. Heermann, S., Spittau, B., Zajzon, K., Schwab, M. H. & Krieglstein, K. Schwann cells migrate along axons in the absence of GDNF signaling. BMC Neurosci. 13, 92 (2012).

22. Furlan, A. & Adameyko, I. Schwann cell precursor: a neural crest cell in disguise? Dev. Biol. 444, S25–S35 (2018).

23. Miyamoto, Y. et al. Neuregulin-1 type III knockout mice exhibit delayed migration of Schwann cell precursors. Biochem. Biophys. Res. Commun. 486, 506–513 (2017).

24. Garratt, A. N., Voiculescu, O., Topilko, P., Charnay, P. & Birchmeier, C. A dual role of erbB2 in myelination and in expansion of the Schwann cell precursor pool. J. Cell Biol. 148, 1035–1046 (2000).

25. Han, Y. et al. Tumor-induced generation of splenic erythroblast-like Ter-cells promotes tumor progression. Cell 173, 634-648.e12 (2018).

26. Malric, L. et al. Inhibiting integrin β8 to differentiate and radiosensitize glioblastoma-initiating cells. Mol. Cancer Res. 17, 384–397 (2019).

27. Su, Z. et al. An investigation of biomarkers derived from legacy microarray data for their utility in the RNA-seq era. Genome Biol. 15, 523 (2014).

28. Molenaar, J. J. et al. Sequencing of neuroblastoma identifies chromothripsis and defects in neuritogenesis genes. Nature 483, 589–93 (2012).

29. Shimada, H. et al. Histopathologic prognostic factors in neuroblastic tumors: definition of subtypes of ganglioneuroblastoma and an age-linked classification of neuroblastomas. J. Natl. Cancer Inst. 73, 405–16 (1984).

30. Ambros, I. M. et al. Role of Ploidy, Chromosome 1p, and Schwann Cells in the Maturation of Neuroblastoma. N. Engl. J. Med. 334, 1505–1511 (1996).

31. Shimada, H. et al. Terminology and morphologic criteria of neuroblastic tumors: recommendations by the International Neuroblastoma Pathology Committee. Cancer 86, 349–63 (1999).

32. Mora, J., Akram, M., Cheung, N. K., Chen, L. & Gerald, W. L. Laser-capture microdissected schwannian and neuroblastic cells in stage 4 neuroblastomas have the same genetic alterations. Med. Pediatr. Oncol. 35, 534–7 (2000).

33. Mora, J. et al. Neuroblastic and Schwannian stromal cells of neuroblastoma are derived from a tumoral progenitor cell. Cancer Res. 61, 6892–6898 (2001).

34. Coco, S. et al. Genome analysis and gene expression profiling of neuroblastoma and ganglioneuroblastoma reveal differences between neuroblastic and Schwannian stromal cells. J. Pathol. 207, 346–357 (2005).

35. Chen, L. et al. CRISPR-Cas9 screen reveals a MYCN-amplified neuroblastoma dependency on EZH2. . Clin. Invest. 128, 446–462 (2018).

36. Wang, C. et al. EZH2 mediates epigenetic silencing of neuroblastoma suppressor genes CASZ1, CLU, RUNX3, and NGFR. Cancer Res. 72, 315–24 (2012).

37. Li, Z. et al. EZH2 regulates neuroblastoma cell differentiation via NTRK1 promoter epigenetic modifications. Oncogene 37, 2714–2727 (2018).

38. Carr-Wilkinson, J. et al. Differentiation of human embryonic stem cells to sympathetic neurons: a potential model for understanding neuroblastoma pathogenesis. Stem Cells Int. 2018, 4391641 (2018).

39. Nakagawara, A. et al. Association between High Levels of Expression of the TRK Gene and Favorable Outcome in Human Neuroblastoma. N. Engl. J. Med. 328, 847–854 (1993).

40. Potzner, M. R. et al. Sequential requirement of Sox4 and Sox11 during development of the sympathetic nervous system. Development 137, 775–84 (2010).

41. Meyers, R. M. et al. Computational correction of copy number effect improves specificity of CRISPR-Cas9 essentiality screens in cancer cells. Nat. Genet. 49, 1779–1784 (2017).

42. La Manno, G. et al. RNA velocity of single cells. Nature 560, 494–498 (2018).

43. Batlle, E. & Clevers, H. Cancer stem cells revisited. Nature Medicine vol. 23 1124–1134 (2017).

44. England, M. A. A colour atlas of life before birth. (Year Book Medical Pub. Inc, 1990).

45. Mayrhofer, M., Viklund, B. & Isaksson, A. Rawcopy: Improved copy number analysis with Affymetrix arrays. Sci. Rep. 6, 36158 (2016).

46. McCarthy, D. J. et al. Cardelino: computational integration of somatic clonal substructure and single-cell transcriptomes. Nat. Methods 17, 414–421 (2020).

