## Supplementary Material: Material/Methods and Supplementary Figures S1-S3 for "Malignant Schwann cell precursors mediate intratumoral plasticity in human neuroblastoma"

**This file includes:**

Materials and Methods
Supplementary figures S1 to S3
References (43 – 45)
Captions for Data S1 to S4

**Other Supplementary Materials for this manuscript include the following:**

Data S1 to S4

### Materials and Methods

Patient material and collection of tumor specimens

Neuroblastoma samples from Swedish patients were collected after parents or guardians had given verbal or written consent. All samples were collected according to permits approved by the Regional Ethics Committee in Stockholm to P.Ko. (reference numbers 03-736, 2009/1369-31/1) and adhered to the Declaration of Helsinki. Clinical data were obtained from hospital records according to the ethical permit. For fetal tissue, an 8 weeks (post-conception) human fetus was retrieved from a clinical routine abortion with written consent from the pregnant woman, after receiving oral and written information. All procedures were performed according to permits from the Ethical Review Authority and the National Board of Health and Welfare. Age was determined by clinical ultrasound assessment, actual crown-rump-length and development of anatomical landmarks *(42)*. The adrenal glands were dissected in DMEM under sterile conditions and stored in ice cold PBS pending analysis.

Dissociation of neuroblastoma into single cells

Fresh tissue obtained from surgery was collected in Media 199 (Gibco) supplemented with 2% (v/v) fetal bovine serum (FBS) on ice. Single cell suspensions of the tumors were obtained by cutting the tumor into small pieces (1mm3) followed by enzymatic dissociation for 45 minutes at 37°C with shaking at 120 rpm using Collagenase I, Collagenase II, Collagenase III, Collagenase IV (all at a concentration of 1 mg/ml; Worthington Biochemical) and Dispase (2 mg/ml; Thermo Fisher) in the presence of DNAse I solution (Thermo Fisher Scientific; 1:100) and RNase Out (Thermo Fisher; 1:1000). Cells were then resuspended in sorting media (Media 199 supplemented with 2% FBS).

Fetal adrenal cell isolation and preparation

Fetal tissue was dissociated using 0.05% Trypsin/0.02% EDTA (Sigma) for 15 minutes at 37°C, with mixing every five minutes. The tissue was then triturated with P-1000 and P-200 pipettes until no cell clumps were visible. The reaction was quenched with PBS containing 10% FBS, cells were then centrifuged, washed twice in PBS, and resuspended in 100 µl PBS with 0,04% bovine serum albumin (BSA; Sigma). Finally, 400 µl ice cold methanol was added for fixation and cells were cryopreserved.

FACS enrichment of viable neuroblastoma cells

Cells from neuroblastoma samples were stained with anti-CD235-PE (Biolegend) and Calcein AM (Thermo Fisher) for 30 minutes at 4°C. Cells were washed twice with Media199 containing 2% FBS followed by DAPI and/or 7AAD staining (1 µg/ml). Flow sorting for live and nonerythroid cells (DAPI/7AAD-negative, Calcein AM-positive, CD235-negative) was performed on a BD FACS Aria III equipped with a 100 µm nozzle (BD Biosciences, San Jose, CA).

Single cell library preparation and sequencing

After FACS sorting, neuroblastoma single-cell libraries were prepared using the Chromium single-cell 3' reagent kit v2 (10x Genomics) according to the manufacturer's recommendations. Libraries were sequenced on the NextSeq 500 platform (Illumina). Fetal adrenal cells were thawed and spun down. Methanol was removed and cells were washed twice in DBPS (Gibco) containing 1% BSA and 0.5 U/µl RNAse OUT. Single-cell libraries were prepared using the Chromium v3 protocol (10x Genomics) and sequenced using the NovaSeq 6000 platform (Illumina).

SNP-array CGH

DNA extraction from neuroblastoma samples was performed using the DNeasy blood and tissue kit (Qiagen) according to the manufacturer's protocol. DNA quality was assessed using the Tapestation platform (Agilent). SNP microarray analysis was performed using the Cytoscan HD Array (Affymetrix) and data was processed using *rawcopy* software *(43)*.

Sectioning of neuroblastoma tumors

Snap frozen neuroblastoma samples were slowly brought from -80°C to +4°C. Samples were fixed for four hours in 4% paraformaldehyde in PBS (pH 7.4) at 4°C with constant rotation, rinsed in PBS three times and cryoprotected by overnight incubation in PBS containing 30% sucrose at 4°C, again with constant rotation. Tissue samples were then embedded in OCT (VWR Chemicals) and frozen at -20°C. Serial sections (10, 14 or 40 µm) were produced from each sample and collected on SuperFrost Plus Adhesion microscope slides (ThermoFisher) and stored at -20°C.

Fluorescent in situ hybridization (FISH)

Cryosections were thawed and dried in room temperature for 30 minutes and rehydrated in PBS for 15 minutes. Sections were then permeabilized in 0.5% Triton X-100 (Thermo Fisher Scientific) in PBS for 20 minutes and incubated in 10 mM sodium citrate buffer (pH 6) for 10 minutes, both at room temperature. After 20 minutes of incubation in 10 mM sodium citrate buffer at 80 °C, slides were cooled to room temperature and washed for five minutes in 2x saline sodium citrate (SSC) buffer. Sections were then incubated in cold 100% acetone at -20°C for five minutes, washed three times in 2x SSC buffer for five minutes each, and subsequently incubated in 50% formamide/2x SSC for four hours at room temperature. 5-TAMRA labelled FISH probes for the PPM1D gene located at chromosomal arm 17q (Empire Genomics) were then diluted 1:5 in FISH probe buffer (Empire Genomics) and applied to the sections. Probes were covered with cover slips, sealed with rubber cement, and pre-hybridized at 37 °C in a dark, moist chamber for 1 hour. After denaturing at 80°C for five minutes, slides were hybridized at 37°C for 48-72 hours. Cover slips were then removed and slides were washed three times in 2x SSC, 37 °C, 5 minutes each. Finally, sections were washed three times in 0.1x SSC at 60°C, 5 minutes each.

Sections were then immunostained according to the protocol described below (without antigen retrieval).

Immunostaining

Slides were brought to room temperature and dried for at least one hour. Antigen retrieval was performed by submerging sections in 1x Target Retrieval Solution (Dako), boiling them briefly and cooling down to room temperature for at least 40 minutes. Sections were washed three times, 10 minutes each, in PBS containing 0.1% Tween-20 (Thermo Fisher Scientific) (PBST), incubated overnight at room temperature with primary antibodies diluted in PBST in a dark, humid chamber. Sections were washed three times, 10 minutes each, in PBST; then incubated with appropriate secondary antibodies and DAPI diluted in PBST at room temperature for 90 minutes in a dark, humid chamber. After incubation, slides were washed three times, 10 minutes each, in PBST and mounted using Mowiol (Merck) mounting medium. The following primary antibodies were used: chicken anti-NF200 (Abcam ab8135; 1:3000), goat anti-SOX10 (Novus Biologicals CL4455; 1:500), SOX10 (R&D Systems AF2864; 1:800), mouse anti-HuC/HuD (Invitrogen A21271, 1:500), rabbit anti-NRG1 (Boster PA1969; 1:500), sheep anti-ERBB3 (R&D Systems AF4518; 1:500), rabbit anti-PDGFRA (CST 5241S; 1:800), mouse anti-ISL1/ISL2 (DSHB AB 2314683. 1:300), chicken anti-MPZ (Abcam ab134439, 1:2000). For detection of primary antibodies, secondary antibodies raised in donkey directed against mouse, rabbit, chicken, sheep, or goat conjugated with Alexa-488, -555 or -647 fluorophores (Thermo Fisher Scientific) were used in 1:1000 dilutions.

Confocal imaging and image analysis

Immunostaining and FISH images were acquired using LSM700 confocal microscope (Zeiss) with 20x, 40x (water) and 63x (oil) objectives. Images were acquired in the .lsm format and processed using Imaris (Bitplane) and ImageJ software. FISH signal was analyzed by evaluating z stacks in 3D mode. All DAPI+ nuclei containing FISH signals of good quality, not obviously cut by the image or section boundaries, were included. The number of FISH signals was evaluated by rotating z stacks, ensuring that FISH signals were located within the DAPI+ nucleus.

Pre-processing of scRNA-seq data

Demultiplexing of .bcl files into .fastq files was performed using *cellranger* 3.0.1 *mkfastq* software (10x Genomics) and alignments to the human (GRCh38) genome reference sequences were performed using *cellranger count*. The reference included all protein coding genes as well as mitochondrial genes for downstream analysis.

All cell barcodes with less than 500 UMIs were excluded. Samples were further filtered one sample at a time where barcodes with percent mitochondrial reads larger than median plus two standard deviations (percent.mito > median + 2sd). Likewise, barcodes with few detected genes were filtered out as $\log_{10}$(nGene) < median - 2sd.

scRNAseq analysis

During the preliminary round of analyses the datasets were aligned using Conos 1.2.1 with $k$ = 15, $k_{self}$= 5 PCA rotation space and angular distance measure. Visualization using largeVis embedding showed a number of continuous bridges connecting the major populations (fig. S1E).

*Bridge tests.* The extent to which the expression of genes in the bridges deviates from simple linear mixing of the connected major subpopulations was tested as follows: expression profile $\boldsymbol{b_i}$ of each cell $i$ in the bridge was modeled using the following linear model: $\boldsymbol{b_i} \sim \boldsymbol{C_1} + \boldsymbol{C_2}$, where $\boldsymbol{C_1}$ and $\boldsymbol{C_2}$ are the cumulative expression profiles of the connected subpopulations (i.e. expression profiles estimated based on a collection of the molecules detected from all of the cells in the cluster, disregarding the cellular barcodes). The modeling was performed with zero intercept. The residual for each gene in the profile $\boldsymbol{b_i}$ was estimated as a ratio of the working (raw) residual and the standard deviation of the gene expression across all of the cells considered in the test (*i.e.* those belonging to the bridge and the two connected populations). Doublet scores were determined using *scrublet* 0.2.1 (fig. S1E).

*Clean alignment.* Cells with scrublet scores above 0.2 were omitted from further analysis. To reduce the impact of cell similarity due to cell cycle patterns, the genes annotated as part of the following GO categories were omitted: GO:0000278, GO:0006260, GO:0007049. *Conos* alignment was then re-constructed using the same parameters as before, and a UMAP embedding was generated (**Fig. 1, B and C**).

*CNV analysis of expression magnitudes (**Fig. 1C**, fig. S1G).* For each sample the regions with genomic gain were determined by log ratio of genomic segments from array CGH data. Mitochondrial genes were omitted. For the initial assessment of the malignant cells (**Fig. 1C, Fig. 2A**), genomic segments with the log2 fold change over 0.15 were selected. For a more stringent evaluation of SCP-like cells (fig. S1G), we also omitted sex chromosomes and required each gained region to cover at least three consecutive probes on the array. The CNV expression score of each cell was then calculated as average expression values of the genes in amplified gained regions. Combining all cells outside of the Adrenergic, Mesenchymal and SCP-like groups into the "other" control group, we then determined a 5% FDR threshold as a minimal score for which the fraction of cells exceeding the threshold contained less than 5% of "other" cells (i.e. 95% of the cells are from the potentially malignant Adrenergic, Mesenchymal and SCP-like populations).

*CNV analysis of allele frequencies.* For each analyzed sample, the overall set of mutations was determined by pseudo-bulk RNAseq data (merged scRNAseq), requiring at least 2 altered alleles and 2 reference alleles. *Cardelino* was used to remap reads to hg19 and call mutations. Alternative and reference allele count matrices were generated for each cell *(44)*. To test for the bias towards non-deleted alleles in the SCP-like and Mesenchymal populations, the following procedure was used: All SNP positions with fewer than 10 minor allele molecule counts were omitted. A subset of the Adrenergic population (called "Adr-", excluding "non-CNV" cluster,

**Fig. 3E**) was used as a gold-standard "malignant" population. All of the SNPs were oriented so that the minor frequency allele within the Adr- population were considered as potentially deleted alleles. To identify SNPs that showed strong statistical evidence for the deletion, we compared the relative allele frequencies within the non-malignant "other" cells, and the Adr-population using Fisher exact test. SNPs with p-value below $10^{-3}$ and minor allele frequency below 0.03 were considered as "definitely deleted" and used in the subsequent step of the analysis. During the next step, the allele pattern of each cell in the sample was used to determine a CNV score, which was estimated as a log likelihood ratio of two models: $H_0$ under which the probability of the cell expressing both alleles was equal to 0.5, and $H_1$ under which the probability of a cell expressing deleted allele was 0.1 and non-deleted 0.9. The likelihood of each model was evaluated using binomial tail probabilities. Occurrence of each allele in a given cell was counted only once (*i.e.* the allele count matrices were binarized for this analysis, to discount mono-allelic bursting effects). To assess whether different subpopulations likely carry malignant cells, resulting allele deletion score distributions of the Adrenergic, Mesenchymal and SCP-like were then compared with the "other" combined control population using one-sided Kolmogorov-Smirnov test (fig. S1H).

*RNA velocity.* To perform RNA velocity analyses, spliced and unspliced reads were recounted using *velocyto* from .bam files of scRNA-seq data *(41)*. We then used *scVelo* (https://github.com/theislab/scvelo) to infer directionality of adrenergic and SCP-like cell transitions. *scVelo* was applied to both merged datasets and individual samples. (**Fig. 3, D and H**, and fig. S3, G and K).

Survival analysis

Bulk RNA-seq data (GSE49711, FPKM) and microarray data (GSE16476; MAS5 signal intensity) were downloaded from GEO. To test if expression of given genes was associated with survival probability, bulk gene expression values from public datasets were stratified into two groups based on gene expression, separating patients with top 25% and bottom 25% expression levels. To test survival association with two genes, bulk samples were separated into four groups with median expression as threshold. A standard Kaplan-Meier survival analysis was then used to determine the association of these groups with survival rate. Kaplan-Meier survival analysis was performed using the *survival* R package. (**Fig. 2F**, fig. S2, E and H, and fig. S3F).

**Figure S1**

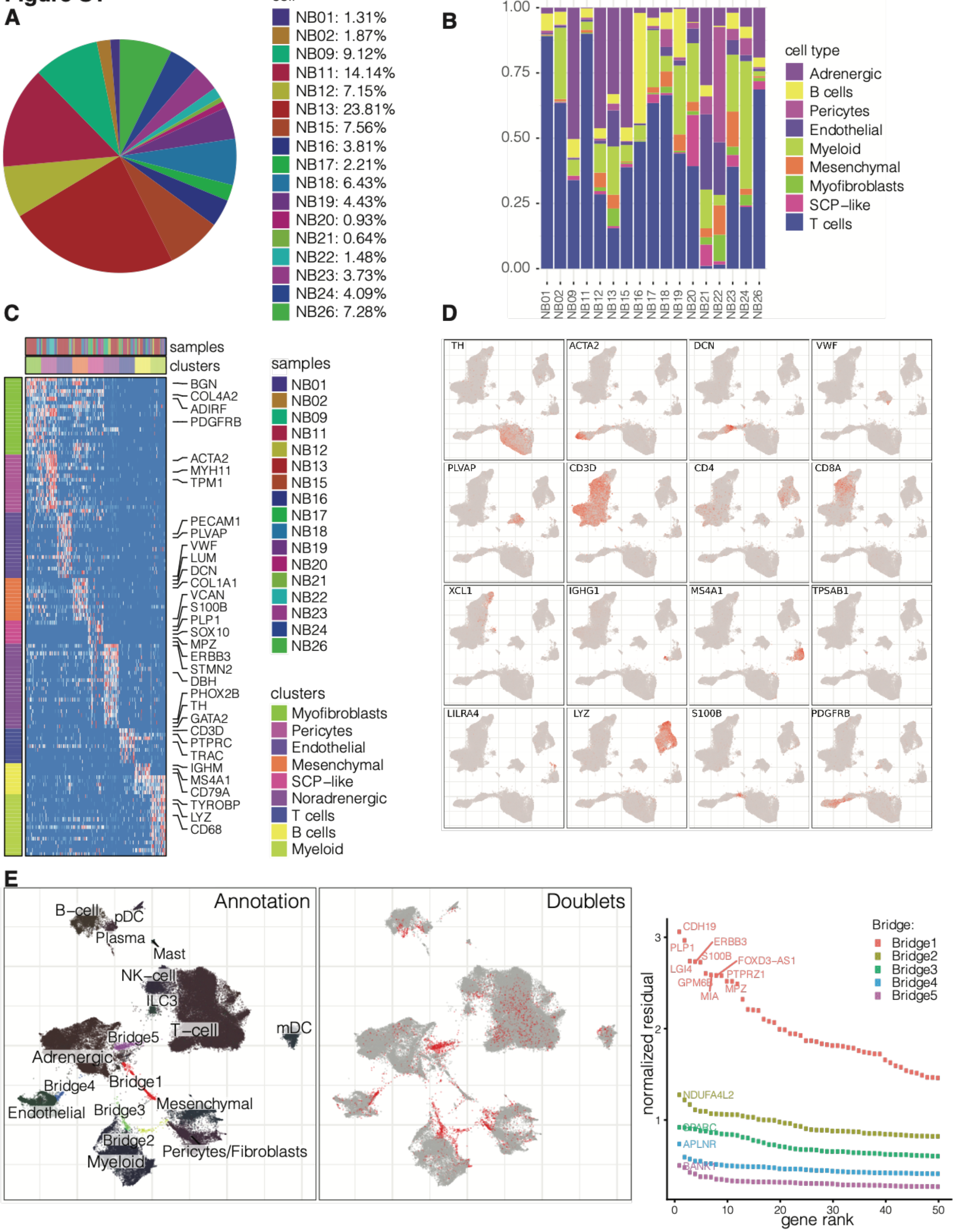
A
cell
NB01: 1.31%
NB02: 1.87%
NB09: 9.12%
NB11: 14.14%
NB12: 7.15%
NB13: 23.81%
NB15: 7.56%
NB16: 3.81%
NB17: 2.21%
NB18: 6.43%
NB19: 4.43%
NB20: 0.93%
NB21: 0.64%
NB22: 1.48%
NB23: 3.73%
NB24: 4.09%
NB26: 7.28%
B
cell type
Adrenergic
B cells
Pericytes
Endothelial
Myeloid
Mesenchymal
Myofibroblasts
SCP−like
T cells
C
samples
clusters
BGN
COL4A2
ADIRF
PDGFRB
ACTA2
MYH11
TPM1
PECAM1
PLVAP
VWF
LUM
DCN
COL1A1
VCAN
S100B
PLP1
SOX10
MPZ
ERBB3
STMN2
DBH
PHOX2B
TH
GATA2
CD3D
PTPRC
TRAC
IGHM
MS4A1
CD79A
TYROBP
LYZ
CD68
clusters
Myofibroblasts
Pericytes
Endothelial
Mesenchymal
SCP−like
Noradrenergic
T cells
B cells
Myeloid
D
TH
ACTA2
DCN
VWF
PLVAP
CD3D
CD4
CD8A
XCL1
IGHG1
MS4A1
TPSAB1
LILRA4
LYZ
S100B
PDGFRB
E
Annotation
Doublets
B−cell
pDC
Plasma
Mast
NK−cell
ILC3
T−cell
mDC
Bridge5
Adrenergic
Bridge4
Bridge1
Endothelial
Bridge3
Mesenchymal
Bridge2
Myeloid
Pericytes/Fibroblasts
normalized residual
gene rank
Bridge:
Bridge1
Bridge2
Bridge3
Bridge4
Bridge5
CDH19
PLP1
ERBB3
S100B
FOXD3−AS1
LGI4
PTPRZ1
GPM6B
MPZ
MIA
NDUFA4L2
SPARC
APLNR
BANK1

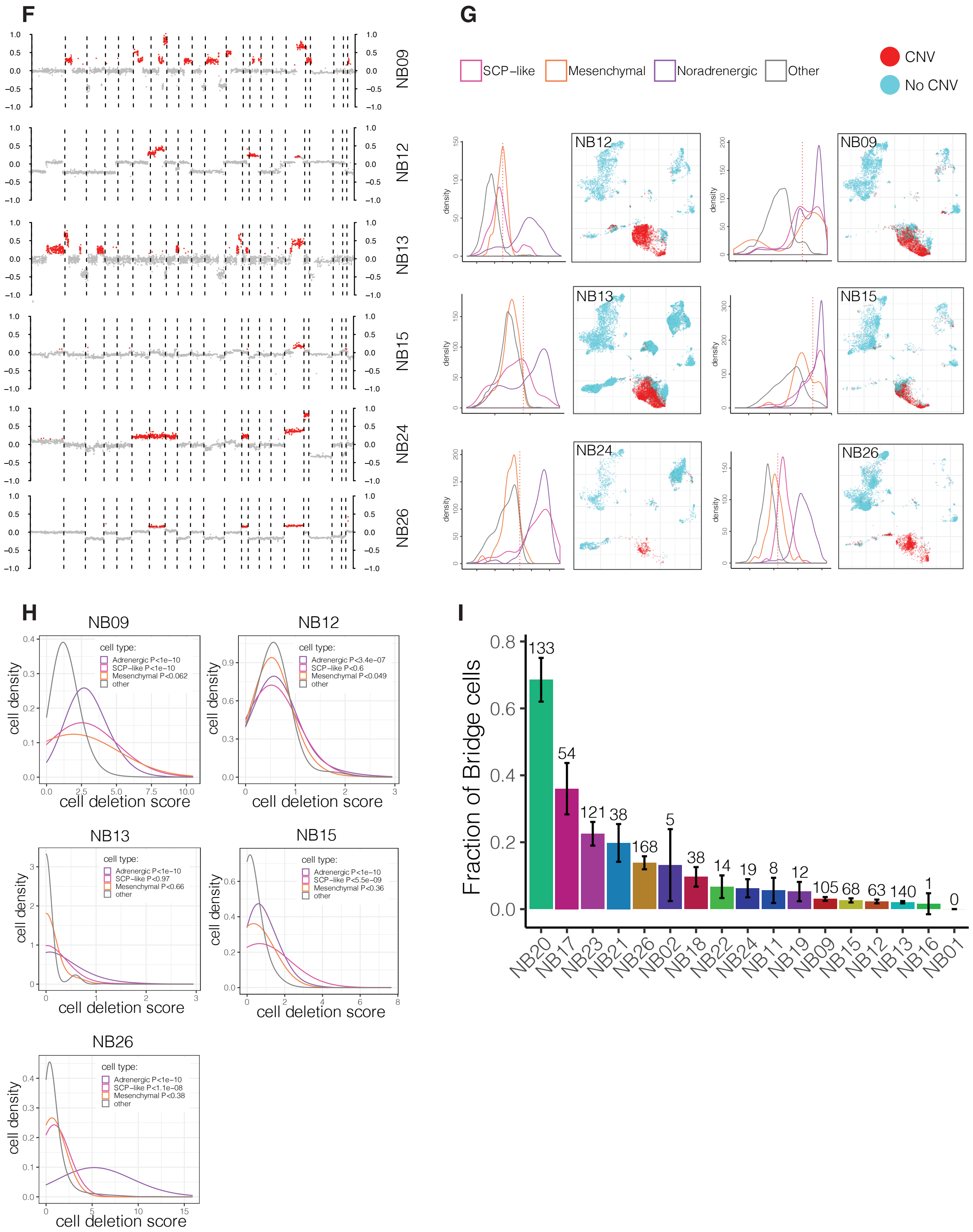

F
NB09
NB12
NB13
NB15
NB24
NB26
G
SCP−like
Mesenchymal
Noradrenergic
Other
CNV
No CNV
NB12
NB09
NB13
NB15
NB24
NB26
density
H
NB09
NB12
NB13
NB15
NB26
cell type:
Adrenergic P<1e−10
SCP−like P<1e−10
Mesenchymal P<0.062
other
Adrenergic P<3.4e−07
SCP−like P<0.6
Mesenchymal P<0.049
other
Adrenergic P<1e−10
SCP−like P<0.97
Mesenchymal P<0.66
other
Adrenergic P<1e−10
SCP−like P<5.5e−09
Mesenchymal P<0.36
other
Adrenergic P<1e−10
SCP−like P<1.1e−08
Mesenchymal P<0.38
other
cell density
cell deletion score
I
Fraction of Bridge cells
133
54
121
38
168
5
38
14
19
8
12
105
68
63
140
1
0
NB20
NB17
NB23
NB21
NB26
NB02
NB18
NB22
NB24
NB11
NB19
NB09
NB15
NB12
NB13
NB16
NB01

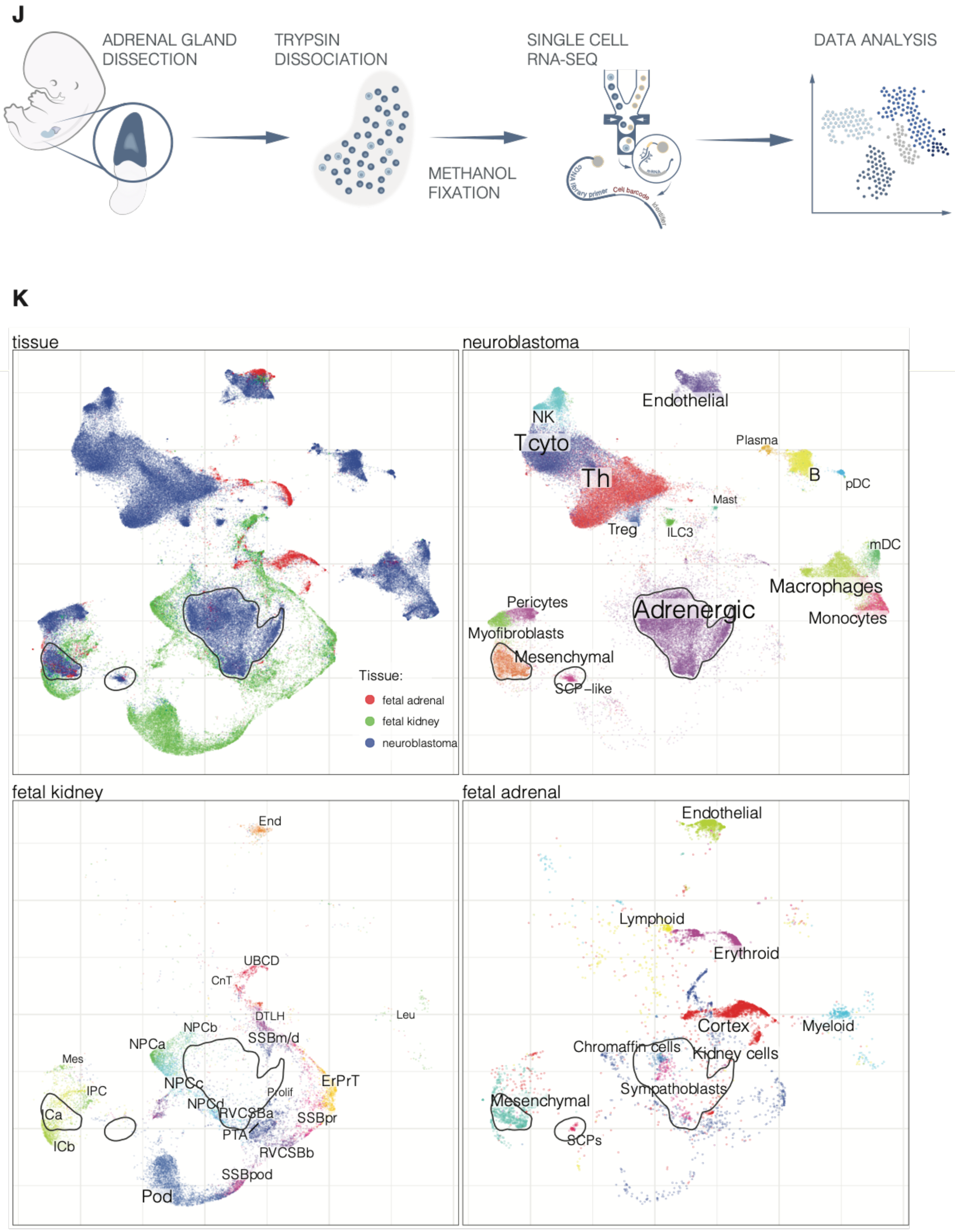
J
ADRENAL GLAND DISSECTION
TRYPSIN DISSOCIATION
METHANOL FIXATION
SINGLE CELL RNA-SEQ
DATA ANALYSIS
K
tissue
Tissue:
fetal adrenal
fetal kidney
neuroblastoma
neuroblastoma
Endothelial
NK
T cyto
Plasma
Th
B
pDC
Mast
Treg
ILC3
mDC
Macrophages
Pericytes
Adrenergic
Monocytes
Myofibroblasts
Mesenchymal
SCP−like
fetal kidney
End
UBCD
CnT
DTLH
Leu
NPCb
SSBm/d
NPCa
Mes
ErPrT
NPCc
IPC
Prolif
NPCd
Ca
RVCSBa
SSBpr
PTA
ICb
RVCSBb
SSBpod
Pod
fetal adrenal
Endothelial
Lymphoid
Erythroid
Cortex
Myeloid
Chromaffin cells
Kidney cells
Sympathoblasts
Mesenchymal
SCPs

**Figure S1. Malignant "bridge" cells in neuroblastoma resemble Schwann cell precursors.** (**A**) Pie chart showing the fraction of cells that each sample contributed to the full dataset. (**B**) Barplot representing the fraction of major cell types within each sample. (**C**) Heatmap showing expression of select marker genes, labelled by cell type and sample. (**D**) Expression of key markers of the major cell types are shown on a joint UMAP embedding (see Fig. 1C). (**E**) Bridge connecting adrenergic and mesenchymal populations shows transient expression features inconsistent with cell doublet capture. (left) Joint analysis of all samples prior to cell filtering is shown as a UMAP embedding. Five bridge populations (Bridge1-5) are shown in bright colors. (middle) Putative doublet cells (with scrublet scores >0.25) are shown in red color. (right) Each bridge was modeled as a linear mixture of the mean expression profiles of the endpoint populations. For each bridge, the plots show mean normalized residuals (y axis) of top 50 genes (x axis) least consistent with a linear mixture. (**F**) Copy number variation (CNV) profiles from array CGH data. Gained genome segments are highlighted (red) for CNV expression score (see Methods). (**G**) (left) Distributions of patient-specific genomic gain expression scores are shown for the potential tumor populations and all other cells in different samples (see Methods). The vertical dashed line shows 5% FDR score cutoff. Only samples with sufficient numbers of CNV genomic gain events are shown. (right) Cells with predicted CNVs are shown in red using the joint UMAP embedding (Fig. 1C). (**H**) Distributions of patient-specific CNV deletion scores are shown for the different cell populations in the samples where sufficient number (>10) of deletion-associated alleles could be identified (see Methods). The statistical significance of the extent to which the deletion score of the different populations exceeds the combined "Other" controls was assessed using one-tailed Kolmogorov-Smirnov test, with the p-value given in the legend of each plot. (**I**) Bar plot showing the abundance of SCP-like bridge cells in each sample, evaluated as a fraction the combined Adrenergic, Mesenchymal and SCP-like populations (y axis). The error bar gives the 95% confidence interval of the proportion estimate for each patient. The numbers above the error bar show the absolute number of the SCP-like cells in the sample. (**J**) Experimental workflow for sampling and processing of normal fetal adrenal tissue. (**K**) Joint alignment of neuroblastoma, fetal adrenal and fetal kidney *(17)* scRNA-seq datasets is shown on a UMAP embedding, colored by data resource (top left), neuroblastoma cell annotation (top right), fetal kidney cell annotation (bottom left) and human adrenal data annotation (bottom right). Solid contours show positions of the neuroblastoma Adrenergic, Mesenchymal and SCP-like populations. Figure 1E of the main manuscript shows a subregion of this alignment.

### Figure S2

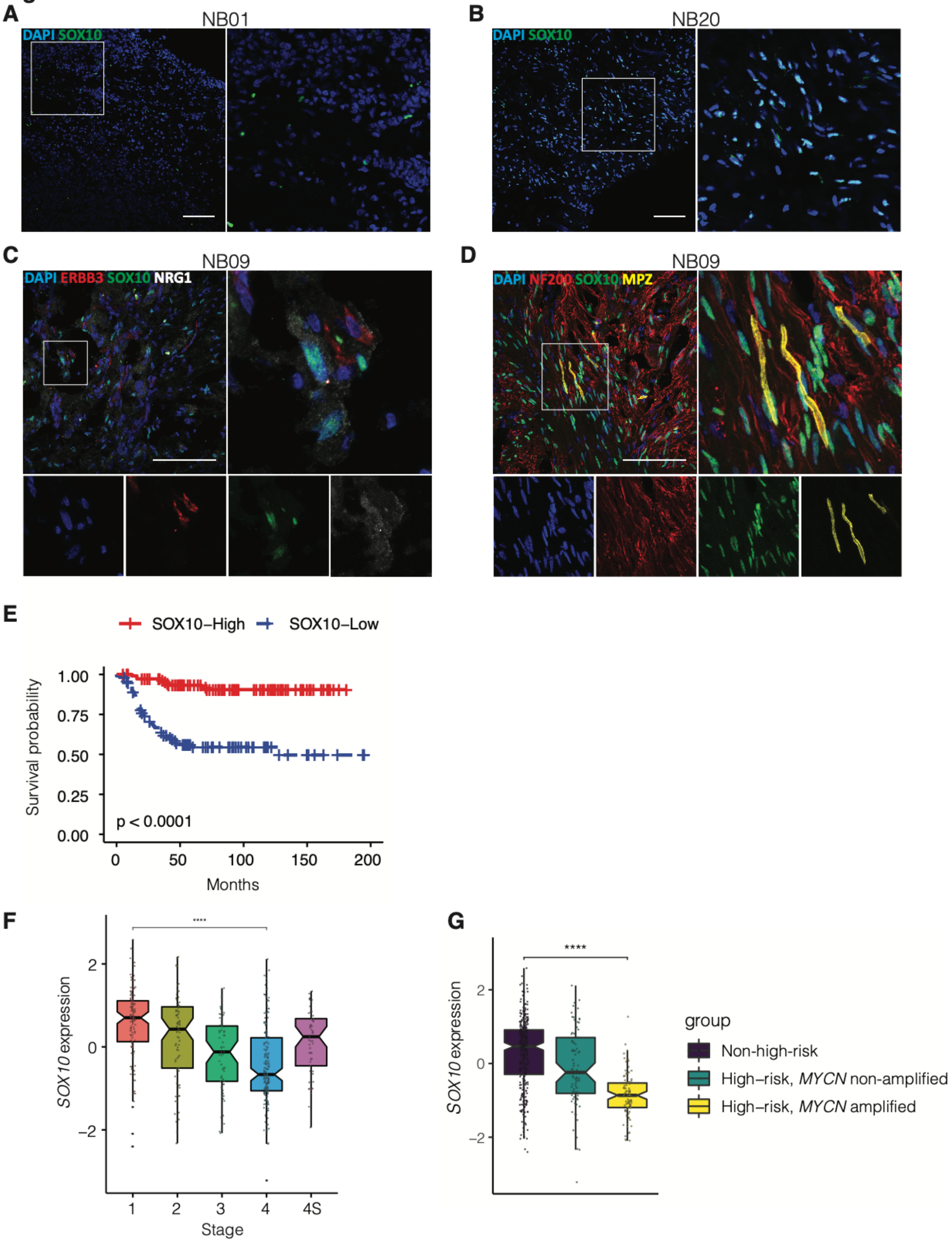

A
NB01
DAPI SOX10
B
NB20
DAPI SOX10
C
NB09
DAPI ERBB3 SOX10 NRG1
D
NB09
DAPI NF200 SOX10 MPZ
E
SOX10−High
SOX10−Low
Survival probability
1.00
0.75
0.50
0.25
0.00
p < 0.0001
0
50
100
150
200
Months
F
SOX10 expression
2
0
−2
****
1
2
3
4
4S
Stage
G
SOX10 expression
2
0
−2
****
group
Non-high-risk
High−risk, MYCN non-amplified
High−risk, MYCN amplified

H

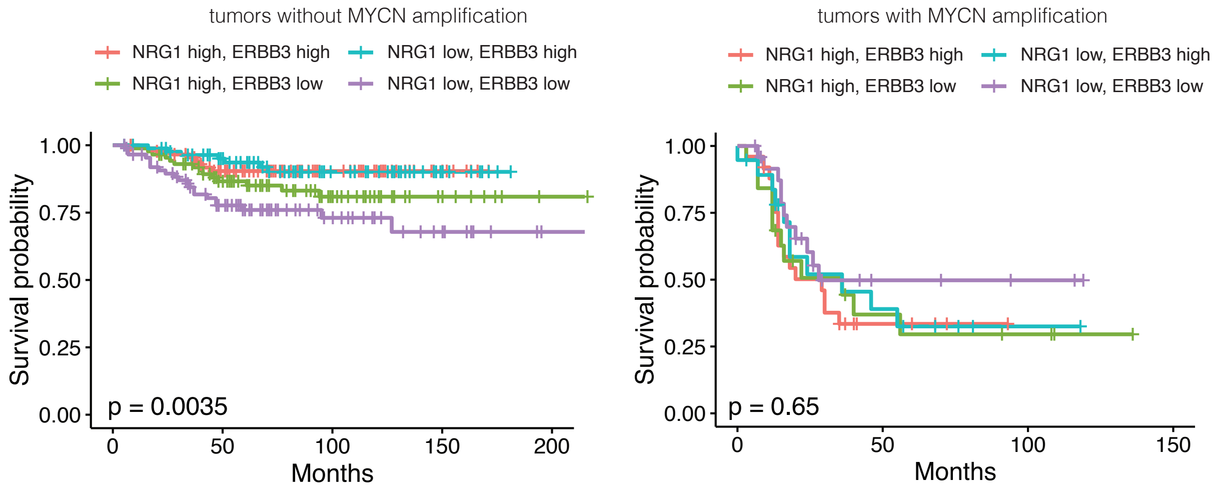


I

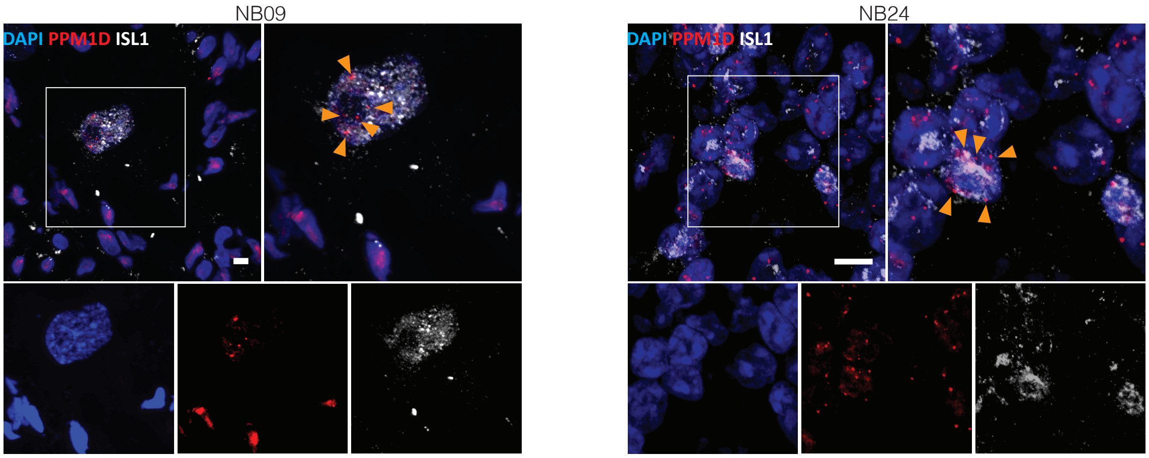


**Figure S2. Interplay between SCP-like neuroblastoma cells and their microenvironment parallel normal development.**
(**A** and **B**) Immunohistochemistry staining for SOX10 (shown in green), demonstrating variations in SOX10+ abundance between samples. The region enlarged in the right panel is marked by the white rectangle in the left panel. (**C**) Immunohistochemistry for SOX10 (green), the receptor tyrosine kinase erbB-3 (ERBB3; shown in red), and its ligand neuregulin-1 (NRG1, shown in white) in sample NB09. Green arrows indicate SOX10+ cells co-expressing ERBB3. White arrows indicate presence of NRG1. Scale bar indicates 100 µm. (**D**) Left panel: Major cell types are labeled on the joint UMAP embedding. Middle and right panel: Expression of NRG1 and ERBB3 in UMAP embedding, illustrating *ERBB3* expression in SCP-like cells and *NRG1* expression in adrenergic cells. (**E**) Immunohistochemistry for neurofilament NF200 (red), SOX10 (green) and myelin protein zero (MPZ; yellow) in sample NB09. Most SOX10+ cells do not express MPZ. Scale bar indicates 100 µm. (**F**) Kaplan–Meier survival curves for neuroblastoma patients (Su et al. dataset, excluding stage 4S, n = 445) according to SOX10

expression status (high or low expression). (**G**) Expression of *SOX10,* Su et al. bulk RNA-seq dataset (all patients, n = 498). Left panel: According to stage. Right panel: According to risk group and *MYCN* status. **** indicates p value < 0.0001. Expression values are shown as log FPKM. (**H**) Kaplan–Meier survival curves for neuroblastoma patients (Su et al. dataset, 4S stage patients excluded, n = 445) are shown for patients stratified into four groups based on the *ERBB3* and *NRG1* expression in *MYCN* non-amplified (left) and *MYCN* amplified (right) samples. (**I**) Validation of adrenergic ISL+ cell malignancy using combined FISH and immunohistochemistry in samples NB24 and NB09. *PPM1D* FISH signal shown in red, ISL1 in white. Scale bar indicates 10 μm. Orange arrows indicate aberrant *PPM1D* signal in ISL1+ cells.

**Figure S3**

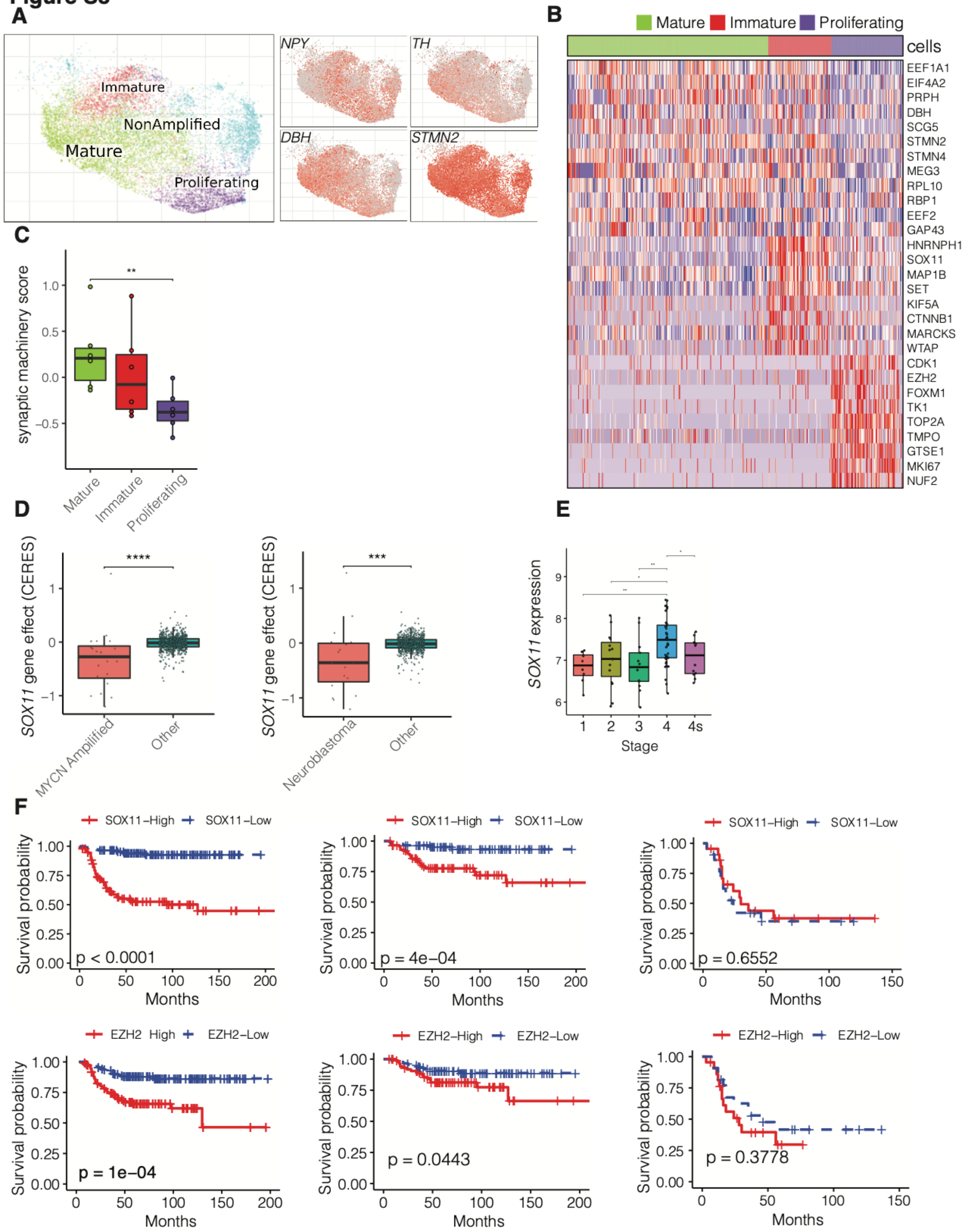
A
Immature
NonAmplified
Mature
Proliferating
NPY
TH
DBH
STMN2
B
Mature
Immature
Proliferating
cells
EEF1A1
EIF4A2
PRPH
DBH
SCG5
STMN2
STMN4
MEG3
RPL10
RBP1
EEF2
GAP43
HNRNPH1
SOX11
MAP1B
SET
KIF5A
CTNNB1
MARCKS
WTAP
CDK1
EZH2
FOXM1
TK1
TOP2A
TMPO
GTSE1
MKI67
NUF2
C
synaptic machinery score
Mature
Immature
Proliferating
D
SOX11 gene effect (CERES)
MYCN Amplified
Other
Neuroblastoma
Other
E
SOX11 expression
Stage
F
SOX11–High
SOX11–Low
Survival probability
Months
p < 0.0001
p = 4e–04
p = 0.6552
EZH2 High
EZH2–High
EZH2–Low
p = 1e–04
p = 0.0443
p = 0.3778

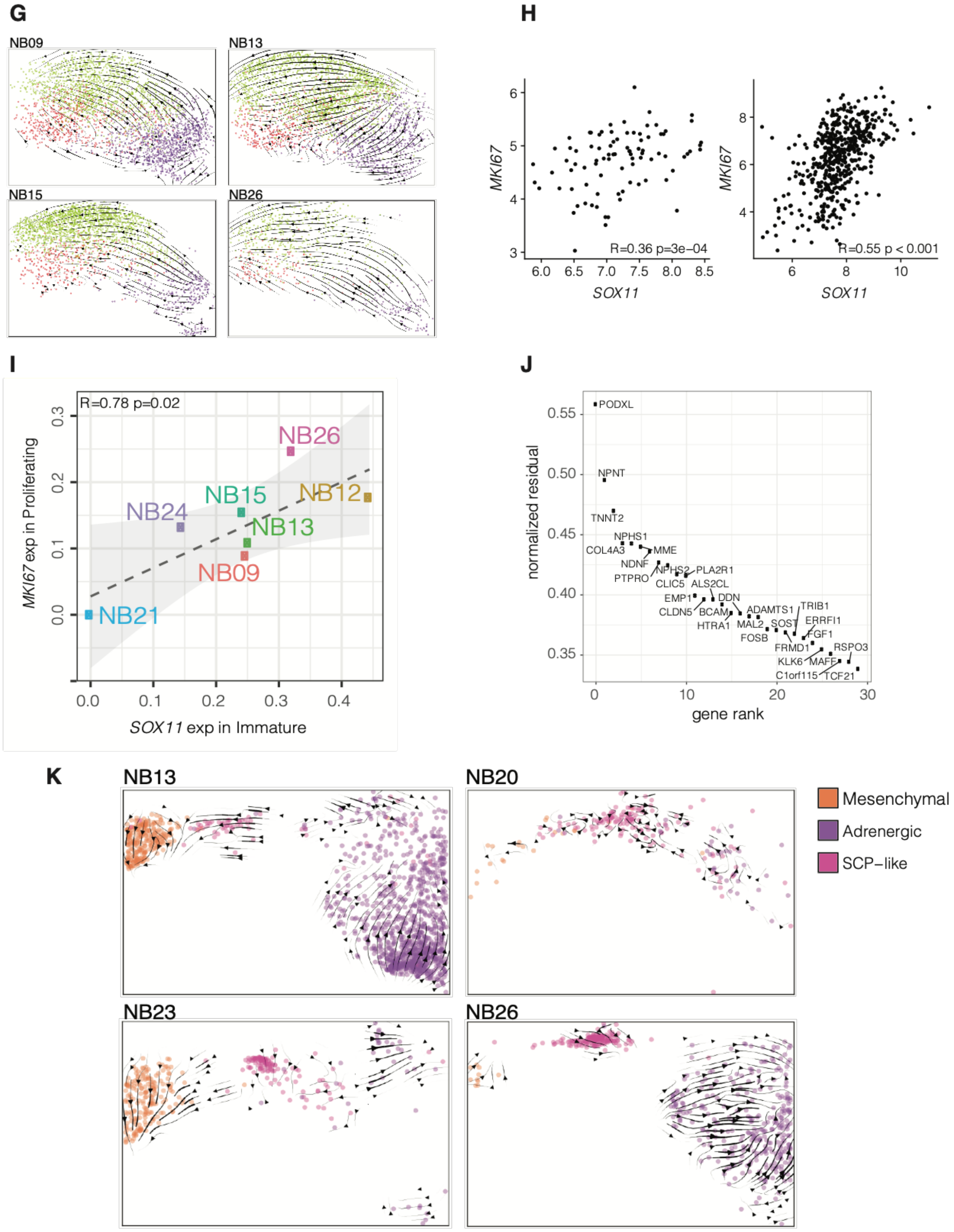


**Figure S3. Adrenergic neuroblastoma cells display varying levels of differentiation.** (**A**) Annotation of adrenergic subpopulations (left) and adrenergic marker genes expression (right) is shown on the corresponding region of the overall UMAP embedding (Fig. 1C). (**B**)

Heatmap showing expression of selected signature genes in *Mature, Immature,* and *Proliferating* adrenergic neuroblastoma cells. (**C**) Average expression (Z scores, see Methods) of genes in the synaptic machinery category (GO:0045202) is shown across different adrenergic subpopulations. Dots represent samples. **: $p < 0.01$. (**D**) Boxplot of CERES scores from CRISPR screens of neuroblastoma cell lines (left) and *MYCN*-amplified cell lines (right), indicating that these are highly dependent on *SOX11*. CERES estimates guide activity scores for genes (Avana dataset, public 20Q1 version). ***: $p < 0.001$. ****: $p < 0.0001$. (**E**) Boxplot of *SOX11* expression according to disease stage (Molenaar et al. dataset, n = 88). *: $p < 0.05$, **: $p < 0.01$. Expression values shown as log MAS5 signal intensity. (**F**) Kaplan–Meier survival curves for neuroblastoma patients (Su et al. dataset, 4S patients excluded, n = 445) according to *SOX11* (top) and *EZH2* (bottom) expression status (high or low expression) in all samples (left), *MYCN* non-amplified samples (middle) and *MYCN*-amplified samples (right). (**G**) RNA velocity analysis showing predicted dynamics within the adrenergic compartment for the individual samples.
(**H**) Scatter plot illustrating significant expression correlation between *SOX11* and *MKI67* in bulk RNA-seq (left panel: Su et al. dataset, n = 498) and microarray (right panel: Molenaar et al. dataset, n = 88) data. Expression values are shown as log FPKM (left panel) and log MAS5 signal intensity (right panel). (**I**) Scatter plot showing significant correlation between *SOX11* expression in *Immature* adrenergic cells and *MKI67* expression in *Proliferating* adrenergic cells.
**(J)** Genes with top 30 residuals deviation from a linear mixture model (see Methods) are shown for the bridge connecting SCP-like and mesenchymal populations in the neuroblastoma samples.
(**K**) RNA velocity analysis of the transitions surrounding SCP-like neuroblastoma state, estimated in individual samples.

**Data S1. (separate file)**
Clinical characteristics of neuroblastoma patients.

**Data S2. (separate file)**
Marker genes used for cell annotations. PubMed IDs for publications supporting the annotation is provided for each gene. The colors correspond to the UMAP embedding shown in Figure 1C.

**Data S3. (separate file)**
Sequencing metrics for all neuroblastoma scRNAseq samples as provided by *cellranger* software.

**Data S4. (separate file)**
Expression of characteristic genes in *Mature, Proliferating,* and *Immature* adrenergic cell populations shown in Fig. 3E and fig. S3B; output from *conos* getDifferentialGenes function. Tables indicate differentially expressed genes, comparing each group against all others using Wilcoxon rank sum test. Z: adjusted Z score; positive values indicate higher expression in a given group. AUC: area under the ROC curve. ExpressionFraction; sum of the gene's expression within the cluster divided by total expression of the gene.
